# Evolution of an extreme hemoglobin phenotype contributed to the sub-Arctic specialization of extinct Steller’s sea cows

**DOI:** 10.1101/2022.08.29.505768

**Authors:** Anthony V. Signore, Phillip R. Morrison, Colin J. Brauner, Angela Fago, Roy E. Weber, Kevin L. Campbell

**Author notes:** National Centre for Foreign Animal Disease, Winnipeg, R3E 3M4, Canada (Anthony V. Signore). Department of Resource Management and Protection, and Biology Department, Vancouver Island University, Nanaimo, V9R 5S5, Canada (Phillip R. Morrison). **Author contributions:** Anthony V Signore, Conceptualization, Project administration, Data curation, Formal analysis, Investigation, Methodology, Visualization, Writing – original draft, Writing – review and editing; Phillip R Morrison, Data curation, Investigation, Methodology, Writing – review and editing; Colin J Brauner, Project administration, Funding acquisition, Resources, Supervision, Writing – review and editing; Angela Fago, Project administration, Funding acquisition, Resources, Supervision, Writing – review and editing; Roy E Weber, Project administration, Funding acquisition, Resources, Supervision, Writing – review and editing; Kevin L Campbell, Conceptualization, Project administration, Funding acquisition, Resources, Formal analysis, Investigation, Methodology, Visualization, Supervision, Writing – original draft, Writing – review and editing. **Competing interests:** Authors declare that they have no competing interests.

## Abstract

The extinct Steller’s sea cow (*Hydrodamalis gigas*; †1768) was a whale-sized marine mammal that manifested profound morphological specializations to exploit the harsh coastal climate of the North Pacific. Yet despite first-hand accounts of their biology, little is known regarding the physiological adjustments underlying their evolution to this environment. Here, the adult-expressed hemoglobin (Hb; α_2_β/δ_2_) of this sirenian is shown to harbor a fixed amino acid replacement at an otherwise invariant position (β/δ82Lys→Asn) that alters multiple aspects of Hb function. First, our functional characterization of recombinant sirenian Hb proteins demonstrate that the Hb–O_2_ affinity of this sub-Arctic species was less affected by temperature than those of living (sub)tropical sea cows. This phenotype presumably safeguarded O_2_ delivery to cool peripheral tissues and largely arises from a reduced intrinsic temperature sensitivity of the *H. gigas* protein. Additional experiments on *H. gigas* β/δ82Asn→Lys mutant Hb further reveal this exchange renders Steller’s sea cow Hb unresponsive to the potent intraerythrocytic allosteric effector 2,3-diphosphoglycerate, a radical modification that is the first documented example of this phenotype among mammals. Notably, β/δ82Lys→Asn moreover underlies the secondary evolution of a reduced blood–O_2_ affinity phenotype that would have promoted heightened tissue and maternal/fetal O_2_ delivery. This conclusion is bolstered by analyses of two Steller’s sea cow prenatal Hb proteins (Hb Gower I; ζ_2_ε_2_ and HbF; α_2_γ_2_) that suggest an exclusive embryonic stage expression pattern, and reveal uncommon replacements in *H. gigas* HbF (γ38Thr→Ile and γ101Glu→Asp) that increased Hb–O_2_ affinity relative to dugong HbF. Finally, the β/δ82Lys→Asn replacement of the adult/fetal protein is shown to increase protein solubility, which may have elevated red blood cell Hb content within both the adult and fetal circulations and contributed to meeting the elevated metabolic (thermoregulatory) requirements and fetal growth rates associated with this species cold adaptation.

## Introduction

The underwater foraging time of mammals is dictated by onboard oxygen stores and the efficiency of their use. Thus, evolutionary increases in oxygen stores, in the form of increased hemoglobin (Hb) and myoglobin—located within erythrocytes and skeletal/cardiac muscle, respectively—are nearly ubiquitous among mammalian divers (Ponganis, 2011). Notable exceptions to this rule are extant sirenians (sea cows), a group of strictly aquatic, (sub)tropical herbivores encompassing only four members; three species of manatee (family Trichechidae) and the dugong, *Dugong dugon* (family Dugongidae). While sirenians are proficient divers, they do not exhibit the greatly elevated body O_2_ stores or an enhanced dive reflex common to other lineages of marine mammals (Blessing, 1972; Scholander and Irving, 1941). Rather, previous work revealed that the sirenian’s secondary transition to aquatic life coincided with a rapid evolution of their Hb encoding genes due, in part, to gene conversion events with a neighboring globin pseudogene (Signore et al., 2019). The resulting high blood–O_2_ affinity phenotype presumably allows extant sea cows to maximize O_2_ extraction from the lungs during submergence at the cost of somewhat reduced O_2_ offloading, thus lowering overall metabolic intensity and fostering a prolonged breath-hold capacity (Signore et al., 2019).

While the relatively limited thermoregulatory capacity of extant sea cows confine them to (sub)tropical waters (Gallivan et al., 1986; Marsh et al., 2011), fossil evidence and first-hand accounts of the sub-Arctic Steller’s sea cow (*Hydrodamalis gigas*) provide insights into the biological and morphological adaptations of this titanic sirenian to the harsh coastal conditions of the North Pacific, where they persisted from the Miocene (5 to 8 million years ago) until their demise in 1768 (Domning, 1976; Heritage and Seiffert, 2022; Stejneger, 1887; Steller, 1751). The retrieval of ancient genetic material from museum specimens has since been instrumental in clarifying the phylogenetic affinities and population history of this species, while providing additional details regarding the evolution of key morphological and physiological attributes (Gaudry et al., 2017; Le Duc et al., 2022; Mirceta et al., 2013; Sharko et al., 2019; Sharko et al., 2021; Signore et al., 2019; Springer et al., 2015) (Fig. 1A). For example, pilot experiments on “resurrected” Steller’s sea cow recombinant Hb demonstrated that the Hb–O_2_ affinity of this lineage secondarily decreased following their divergence from dugongs between the mid Oligocene and early Miocene (Signore et al., 2019). While sirenians do not possess the capacity for non-shivering thermogenesis due to pseudogenization of the *UCP1* gene (Gaudry et al., 2017), the reduced Hb–O_2_ affinity shift in Steller’s sea cow Hb was speculated to have promoted increased O_2_ offloading to fuel increased thermogenesis to help cope with exposure to cold sub-Arctic waters. Although the Hb of this species accumulated 11 amino acid replacements since its divergence from the dugong (Figs. S1, S2), this decrease in Hb–O_2_ affinity was hypothesized to arise from a highly unusual 82Lys→Asn exchange in the chimeric β-type (β/δ) chain (Signore et al., 2019). Data mined from more recent ancient DNA studies (Le Duc et al., 2022; Sharko et al., 2021) confirms that this substitution was fixed in the last remaining Steller’s sea cow population (Fig. 1B), though the specific functional effect(s) of this substitution have not been characterized. This replacement is intriguing not only because β82Lys is invariant among characterized mammalian Hbs, but because human variants with substitutions at this position display profound alterations in both structural and functional properties (Abraham et al., 2011; Bonaventura et al., 1976; Ikkala et al., 1976; Sugihara et al., 1985). For example, the human Hb Providence (β82Lys→Asn) variant exhibits a decreased inherent Hb–O_2_ affinity and markedly reduced sensitivity to the allosteric effectors 2,3-diphosphoglycerate (DPG), Cl^-^, and H^+^ (Abraham et al., 2011; Bardakjian et al., 1985; Bonaventura et al., 1976; Charache et al., 1977; Weickert et al., 1999), all of which preferentially bind and stabilize the (low O_2_ affinity) deoxy-state conformation of the protein. However, opposite to what was suggested for Steller’s sea cows (Signore et al., 2019), this exchange causes the whole blood O_2_ affinity of Hb Providence carriers to be noticeably higher than that of the general population (Bardakjian et al., 1985; Weber and Campbell, 2011). It is thus unclear if and how the Steller’s sea cow β/δ82Lys→Asn replacement underlies the lower Hb–O_2_ affinity of this extinct species relative to other sirenians, or whether this attribute arises from one (or more) of the other 10 residue replacements that evolved in this lineage.

**Figure 1.**
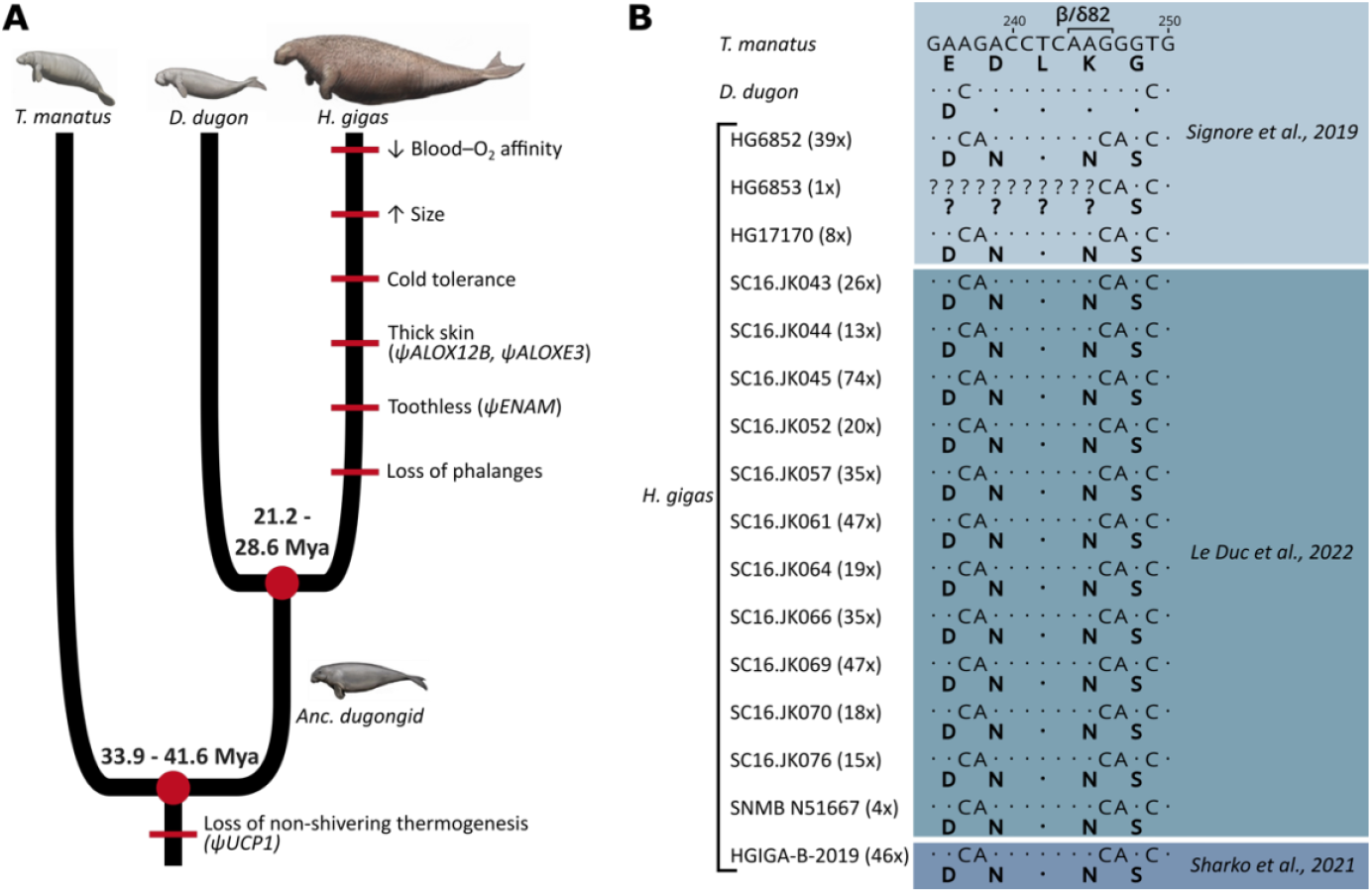
Evolution of notable morphological and genetic attributes within Sirenia. A) Phenotypic innovations contributing to the unique biology of Steller’s sea cows are mapped along the sirenian phylogeny (red bars) with the underlying genetic causes shown in brackets where known. Note that the red bars do not represent the dating of these traits and that their placement order is arbitrary. Divergence dates in millions of years (Mya) are based on Springer et al. (2015) and Heritage and Seiffert (2022). The ancestral dugongid (Anc. dugongid) is represented by a late-Oligocene *Metaxytherium spp*. Images by Carl Buell and are used with permission. B) Partial nucleotide alignment of the sirenian β/δ-globin gene encoding the central region of the 2,3-diphosphoglycerate binding pocket of hemoglobin; corresponding amino acid residues (bolded) are provided below each sequence. The G→C nucleotide mutation underlying the otherwise invariant amino acid substitution (β/δ82Lys→Asn; K→N) of Steller’s sea cow (*Hydrodamalis gigas*) hemoglobin is apparent in all 16 individuals for which sequence data is available. Values in brackets next to each *H. gigas* specimen represent the depth of sequence coverage for this nucleotide. Dots represent sequence identity with the Florida manatee (*Trichechus manatus*).

The β/δ82Lys→Asn exchange also raises other evolutionary significant questions, as it is predicted to have altered multiple aspects of Hb function that may lead to antagonistic pleiotropic effects. Notably, this exchange may detrimentally increase the effect of temperature on Hb–O_2_ binding and release (Signore et al., 2019). The formation of the weak covalent bond between O_2_ and the heme iron requires free energy, thus dictating an inverse relationship between Hb–O_2_ affinity and temperature (Weber and Campbell, 2011). In temperate and Arctic endotherms this inherent attribute of Hb potentially impedes O_2_ delivery to the limbs and flukes, which are maintained at substantively lower temperatures to minimize heat loss and hence energy requirements (Campbell and Hofreiter, 2015). Accordingly, heterothermic mammals generally possess Hbs whose O_2_ binding properties are less sensitive to temperature than the Hbs of non-cold-adapted species, thereby maintaining sufficient O_2_ offloading in the face of decreasing tissue temperatures. This reduction in thermal sensitivity (quantified as the overall enthalpy of oxygenation, ΔH’), appears to predominantly arise from an increased interaction between allosteric effectors and the Hb moiety (an exothermic process), which releases additional heat to assist with deoxygenation (Weber and Campbell, 2011). Hence, the *H. gigas* β/δ82Lys→Asn replacement, which deletes integral binding sites for the heterotropic ligands Cl^-^ and DPG (Bonaventura et al., 1976), is puzzling in that it is expected to maladaptively increase the effect of temperature on O_2_ uptake and release in the blood of the sub-Arctic Steller’s sea cow.

Taken together, it remains unknown whether the *H. gigas* β/δ82Lys→Asn residue exchange contributed adaptively to the species biology or is instead linked to small population sizes (e.g., genetic drift) over the past half million years (Le Duc et al., 2022; Sharko et al., 2021). To unravel the combined effects of evolved amino acid replacements on hemoglobin function in relation to the extreme thermal biology of the extinct Steller’s sea cow, we synthesized recombinant Hb proteins of this extinct species together with those of the extant dugong (*Dugong dugon*) and Florida manatee (*Trichechus manatus latirostris*), and measured their O_2_ binding properties, relative solubility, responses to allosteric effectors, and thermal sensitivities. We also synthesized a *H. gigas* β/δ82Asn→Lys Hb mutant to assess the specific effects of this exchange, together with the Hb of the last common ancestor (‘ancestral dugongid’) shared between the dugong and Steller’s sea cow (Fig. 1A) in order to assess the directionality of evolved physicochemical changes in Hb function.

## Results and Discussion

### O_2_ Affinity of Sirenian Hbs

Measured O_2_-equilibrium curves of the five examined Hbs revealed marked differences in intrinsic O_2_ affinity (Figs. 2A, S3, S4 and Table S1). In the absence of allosteric effectors (pH 7.2, 37°C), the P_50_ (the O_2_ tension resulting in 50% Hb–O_2_ saturation) of Steller’s sea cow Hb (P_50_ = 8.8 mm Hg) is ∼2 times higher than that of dugong (3.5 mm Hg), ancestral dugongid (4.3 mm Hg), and manatee (5.4 mm Hg) Hbs under the same conditions (Figs. 2A, S3 and Table S1). Site directed mutagenesis experiments reveal that the increased intrinsic P_50_ of Steller’s sea cow Hb predominantly arises from the β/δ82Asn substitution, as the β/δ82Asn→Lys mutant exhibits an intrinsic P_50_ similar to that of the ancestral dugongid (Fig. 2A, S4). Of note, the O_2_ affinity of dugong, ancestral dugongid, and manatee Hbs was reduced in the presence of Cl^-^ and DPG (P_50_ = 10.2, 9.9, and 10.9 mm Hg, respectively) by a similar degree to that of Asian elephant Hb (Campbell et al., 2010b). This finding extends previous studies conducted on sirenian Hbs (Farmer et al., 1979; McCabe et al., 1978; Signore et al., 2019), and reveals that the high O_2_ affinity of dugong and manatee blood is not attributable to decreased allosteric effector sensitivity. Conversely, Steller’s sea cow Hb was shown to be markedly less responsive to allosteric effectors, as only a moderate reduction in O_2_–affinity was observed in the presence of Cl^-^ and DPG (P_50_ = 14.3 mm Hg). When the effects of these allosteric effectors were measured individually, Steller’s sea cow Hb exhibits lower DPG, Cl^-^, and H^+^ (Bohr) effects relative to those of the ancestral dugongid and β/δ82Asn→Lys mutant (Fig. 2C-E; Table S1). These data confirm that a high intrinsic (i.e. in the absence of allosteric effectors) Hb–O_2_ affinity is an ancient sirenian trait that likely aided the transition of the group to the aquatic environment, and that Hb–O_2_ (and hence whole blood) affinity was secondarily reduced in the Steller’s sea cow lineage (Signore et al., 2019). This latter finding contrasts with allometric expectations for mammals—whereby blood O_2_ affinity and body mass are inversely correlated (Schmidt-Neilsen and Larimer, 1958)—and thus further suggests this modification served an adaptive function in this extinct species.

**Figure 2.**
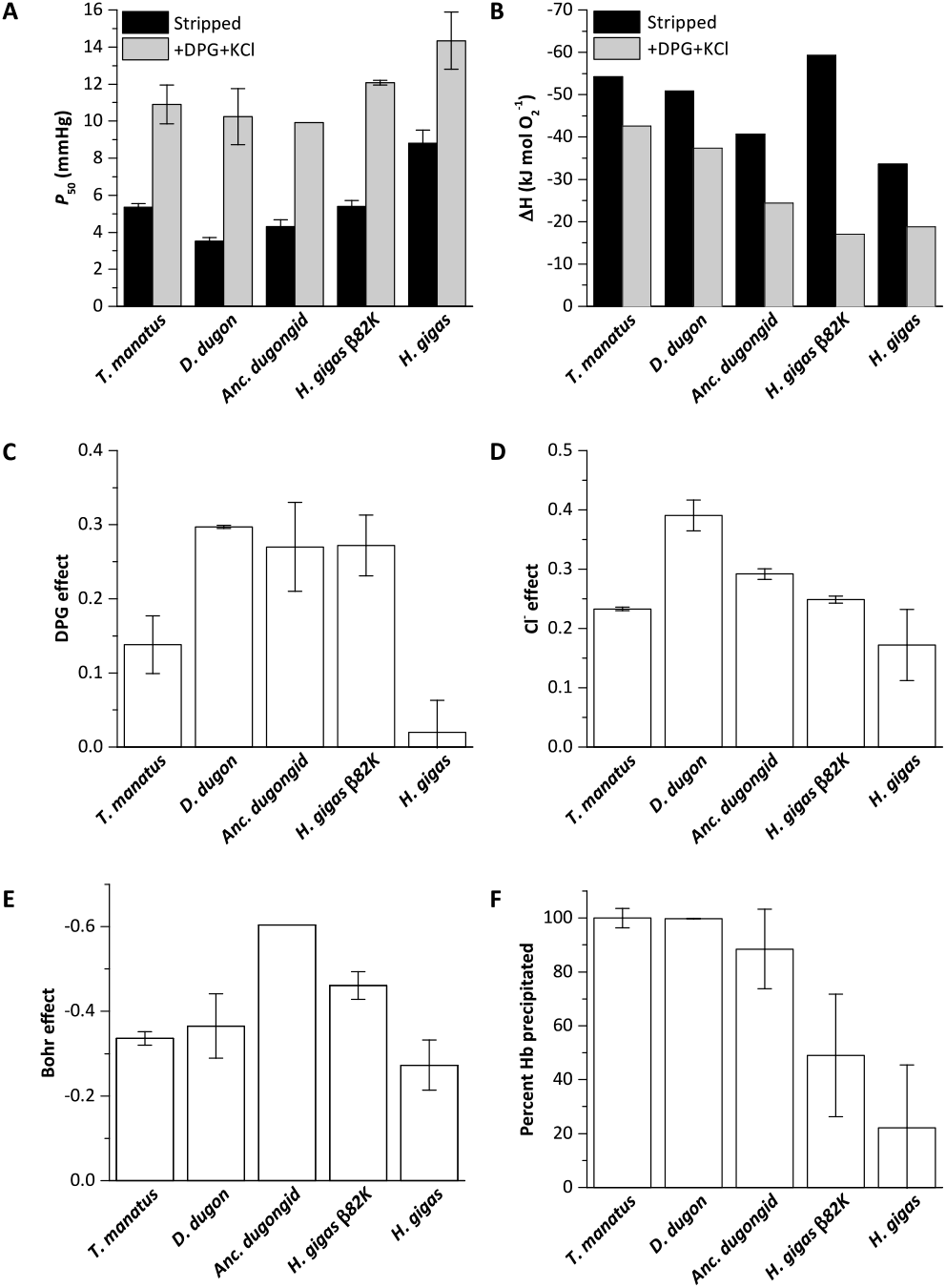
Biochemical properties of hemoglobins (Hbs) from manatee (*Trichechus manatus*), dugong (*Dugong dugon*), ancestral dugongid (Anc. dugongid), Steller’s sea cow β/δ82Asn→Lys mutant (*H. gigas* β82K), and wild-type Steller’s sea cow (*Hydrodamalis gigas*). All values were measured at 37°C and corrected to pH 7.2 (±SE of the regression estimate). A) Oxygen tensions at half O_2_ saturation (P_50_) in the absence (stripped) and presence of allosteric cofactors (2-fold molar excess of 2,3-diphosphoglycerate (DPG) and 0.1 M KCl). B) The enthalpy of oxygenation (ΔH) between 25 and 37°C in stripped Hb and in the presence of allosteric cofactors (2-fold molar excess DPG and 0.1 M KCl). C) The effect of DPG on sirenian Hbs determined from logP_50_^(0.5 mM DPG)^ – logP_50_^(stripped)^. D) The effect of chloride on sirenian Hbs determined from logP_50_^(0.1 M KCl)^ –logP_50_^(stripped)^. E) The Bohr effect of sirenian Hbs in the presence of allosteric cofactors (2-fold molar excess DPG and 0.1 M KCl), as calculated from ΔlogP_50_/ΔpH over the pH range 6.9 and 7.8. F). The relative solubility of sirenian Hbs is denoted by the percentage of Hb protein precipitated by the addition of 3 M ammonium sulfate.

The single most distinct feature of *H. gigas* Hb is the lack of a discernable effect of DPG on P_50_ (ΔlogP_50_^(DPG – stripped)^ = 0.02 at 37°C and pH 7.2; Fig. 2C and Table S1), relative to the Hbs of the ancestral dugongid (0.27) and the extant manatee and dugong (0.14 and 0.30, respectively). This intracellular effector generally occurs in equimolar concentrations to Hb (Bunn, 1980) and strongly decreases the O_2_ affinity of most mammalian Hbs via direct electrostatic interactions with β2His and β82Lys (Fig. 3A), together with potential water-mediated interactions with β143His and the α-NH_2_ group of 1Val of the β_2_ chain (Richard et al., 1993). However, unlike the other (ionizable) residues whose ability to interact with DPG is highly pH dependent, β82Lys is strongly cationic and thus is able to bind DPG across the entire physiological pH range. Presumably arising from this indispensable role in DPG binding, this residue is uniformly conserved in mammalian Hbs (Fig. 3A), with the exception of several heterozygous adult human HbA carriers with substitutions at this position (Ikkala et al., 1976; Moo-Penn et al., 1976; Sugihara et al., 1985). Given that none of the other six β/δ-chain replacements that evolved on the Steller’s sea cow branch (Figs. 3B, S1 and S2) are implicated in DPG binding, the deletion of the integral DPG binding site at β/δ82 in Steller’s sea cow Hb is fully consistent with its inability to bind DPG (Fig. 3B). This conclusion is further supported by our measurements on the Steller’s sea cow β/δ82Asn→Lys mutant, which show that reversion to the ancestral state restores the DPG effect to the same level observed in ancestral dugongid Hb (Fig. 2C). Notably, and despite possessing the identical DPG binding site residues as Hb Providence, the *H. gigas* protein exhibits a distinctly lower DPG effect than this human variant (0.08; 17,20). The lower DPG sensitivity of Steller’s sea cow Hb thus implicates an epistatic contribution from other amino acids in the vicinity of the DPG pocket. Importantly, the Lys→Asn replacement in the DPG binding pocket causes the O_2_ affinity of human HbA to increase in the presence of allosteric cofactors (20,23), whereas results presented in Fig. 2A show Steller’s sea cow Hb–O_2_ affinity is reduced relative to its ancestors carrying β/δ82Lys under all test conditions. This observation highlights a growing body of research indicating that both the direction and overall phenotypic effect of specific amino acid substitutions may be conditional on the genetic background in which they occur (Natarajan et al., 2023; Storz, 2016).

**Figure 3.**
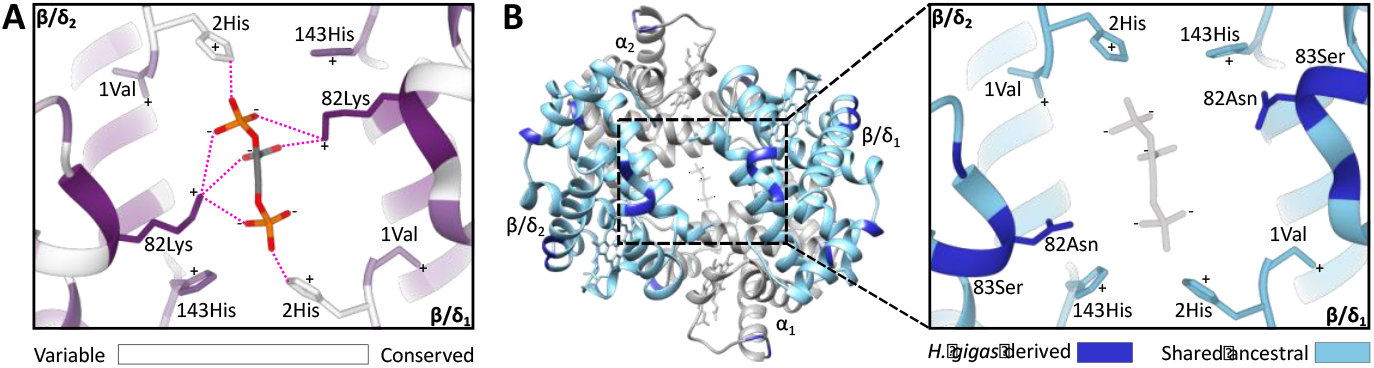
Homology models of the DPG binding site in ancestral dugongid and Steller’s sea cow (*Hydrodamalis gigas*) hemoglobin. A) Model of the ancestral dugongid DPG binding site. Amino acids are colored according to the degree of sequence conservation. Notably, β/δ82Lys shows the highest level of sequence conservation, as it is able to bind to multiple sites on the DPG molecule (indicated by dashed pink lines), whereas β/δ2His is only able to directly interact with DPG in the protonated state. B) Model of Steller’s sea cow hemoglobin (left) and a close-up of the DPG binding pocket (right). Dark blue colored residues represent the 11 *H. gigas* specific substitutions, while those in light blue denote the ancestral state. Homology modelling illustrates how the replacement of β/δ82Lys with neutral Asn inhibits DPG binding to the hemoglobin molecule.

The insensitivity of *H. gigas* Hb to DPG is also notable as it would have markedly reduced their capacity to modulate blood–O_2_ affinity *in vivo* (e.g., seasonally), and is the first demonstrated example of a genuine DPG insensitive Hb phenotype among mammals. While eastern moles (*Scalopus aquaticus*) are a possible exception (Campbell et al., 2010a), feliformid carnivores, ruminants, and two species of lemurs have also traditionally been placed in the ‘DPG insensitive’ category (Bunn, 1980) despite the fact their Hbs are moderately responsive to DPG in the absence of Cl^-^ (Bonaventura et al., 1976; Fronticelli et al., 1988; Janecka et al., 2015; Perutz et al., 1993). Nonetheless, red blood cell DPG concentrations of species with suppressed DPG sensitivities are markedly reduced relative to mammals whose Hb–O_2_ affinity is regulated by DPG (<0.1-1.0 mM vs. 4-10 mM, respectively; (Bunn, 1980)). While the potential benefits of a low DPG sensitivity phenotype have been debated (Campbell et al., 2010a; Kay, 1977), approximately 20% of glucose uptake by human erythrocytes is diverted to DPG synthesis via the Rapport-Luebering shunt (Duhm et al., 1968), thereby bypassing production of both ATP molecules generated via the anaerobic substrate level phosphorylation pathway (Bunn, 1980; Kauffman et al., 2002; Rapoport and Luebering, 1950). Accordingly, since each molecule of DPG produced comes at the expense of producing an ATP molecule, the probability that Steller’s sea cow blood similarly contained low levels of this organophosphate is high.

### Hb Solubility

Ectopic expression of human Hb Providence β82Asp mutants in *E. coli* has been shown to increase soluble protein production by 47-116% relative to the expression of human Hb variants not carrying this substitution (Weickert et al., 1999). Consistent with this observation, we found that Steller’s sea cow Hb is more soluble than those of other sirenians and the engineered *H. gigas* β/δ82Asn→Lys mutant (Fig. 2F and S5). While the precise mechanism underlying this phenomenon is unknown, Hb Providence variants exhibit sharp reductions in irreversible oxidative damage of nearby β93Cys that initiates Hb denaturation (Abraham et al., 2011; Jana et al., 2020; Strader et al., 2017). These β82 replacements thereby presumably decrease the rate of Hb turnover and increase the half-life of the protein (Strader et al., 2017), which may contribute to the mild polycythemia in humans carrying this substitution (Bardakjian et al., 1985; Moo-Penn et al., 1976). Blood with an elevated O_2_ carrying capacity is typical of most mammalian divers, where it increases onboard O_2_ stores and extends dive times (Ponganis, 2011), but is not observed in extant sirenians (Farmer et al., 1979; White et al., 1976; Wong et al., 2018). However, any solubility driven increases in red blood cell Hb concentration resulting from the β82Lys→Asn exchange would have allowed Steller’s sea cows to maintain an elevated rate of tissue O_2_ delivery to meet their metabolic demands during extended underwater foraging intervals. Although this species was presumably unable to completely submerge (Domning, 2022; Steller, 1751), this conjecture is corroborated by Steller’s account that, “*they keep their heads always under water* [foraging], *without regard to life and safety*” (Steller, 1751).

### Thermal Sensitivity

The invariant energy change associated with forming the weak covalent bond between O_2_ and the heme iron (i.e. the enthalpy of heme oxygenation; ΔH^O2^) is exothermic (−59 kJ mol^-1^ O_2_) (Atha and Ackers, 1974), and only moderately opposed by the endothermic solubilization of O_2_ (ΔH^H2O^; 12.55 kJ mol^-1^ O_2_), resulting in an inverse relationship between temperature and Hb– O_2_ affinity. However, the heat of the T→R conformational change (ΔH^T→R^), and the oxygenation-linked dissociation of H^+^ (ΔH^H+^), Cl^-^ (ΔH^Cl-^), and DPG (ΔH^DPG^) may offset this relationship, such that the overall enthalpy of Hb oxygenation (ΔH’) can become greatly minimized or even endothermic (Weber and Campbell, 2011; Weber et al., 2010). By facilitating adequate oxygenation of cool peripheral tissues, Hbs with numerically low ΔH’ values are interpreted to be adaptive for cold-tolerant, regionally heterothermic mammals. The evolution of this phenotype has predominantly been attributed to the formation of additional heterotropic ligand binding sites on the protein moiety, as has previously been demonstrated for the woolly mammoth, *Mammuthus primigenius* (Campbell et al., 2010b; Weber and Campbell, 2011). Conversely, Steller’s sea cow Hb lacks heterotropic binding of DPG and displays lower Bohr (H^+^) and Cl^-^ effects than all other sirenian Hbs measured (Fig. 2; Table S1), yet exhibits a ΔH’ value (−18.8 kJ mol^-1^ O_2_; Fig. 2B) that is close to that of mammoth Hb (−17.2 kJ mol^-1^ O_2_) (Campbell et al., 2010b). This striking convergence largely arises from the inherently low ΔH of stripped Steller’s sea cow Hb (−34.2 kJ mol^-1^ O_2_) relative to dugong, ancestral dugongid, and manatee Hbs (range: -50.1 to -58.6 kJ mol^-1^ O_2_) at pH 7.8—where oxygenation-linked binding of protons is minimal—and indicates that structural differences modifying the T→R transition largely underlie the low thermal sensitivity of *H. gigas* Hb. Recent studies have shown that a large positive ΔH^T→R^ may similarly contribute to the low ΔH’ of deer mouse, cow, shrew, and mole Hbs (Campbell et al., 2010a; Campbell et al., 2012; Jensen et al., 2016; Signore et al., 2012; Weber et al., 2014) suggesting that this potential mechanism of temperature adaptation may be more widespread than previously appreciated. Our experiments with the Steller’s sea cow β82Asn→Lys mutant further implicate substitutions at this position as a key factor underlying the inherently low ΔH of the protein, as this modified protein displays a greatly increased ΔH in the absence of allosteric effectors relative to the wild-type *Hydrodamalis* protein (Fig. 2B). Interestingly, despite these inherent ΔH differences between the mutant and wild-type Steller’s sea cow Hbs, their ΔH’ values are indistinguishable in the presence of allosteric effectors (Fig. 2B). These data suggest that β82Asn uncouples thermal sensitivity from DPG concentration, permanently conferring the *H. gigas* protein with a numerically low ΔH’ by genetic assimilation while simultaneously eliminating the energetic cost of DPG production within the red blood cells.

Given both the marked functional changes observed for *H. gigas* Hb and the correspondingly large ecological and thermal shifts encountered by ancestral hydrodamalines following their exploitation of the North Pacific in the Miocene (Heritage and Seiffert, 2022), it is surprising that previous work failed to provide evidence for positive selection or an accelerated amino acid substitution rate for any globin loci in the Steller’s sea cow branch (Signore et al., 2019). However, this result is not unique to hydrodamalines, as the Hb coding genes of woolly mammoths and stem penguins also lack statistically significant signatures of positive selection accompanying their niche transitions despite clear directional changes in their Hb properties (Campbell et al., 2010b; Signore et al., 2021).

### Paleophysiology of Steller’s sea cows

The posthumously published behavioral and anatomical accounts of the last remaining *H. gigas* population by naturalist Georg Wilhelm Steller while stranded on Bering Island (55°N, 166°E) in 1741/1742 provide a rich tapestry to interpret the paleophysiology of this colossal marine herbivore. For example, their protective thick bark-like hide and extensive blubber layer give credence to the extreme nature of their shallow rocky and (during winter) ice strewn habitat (Le Duc et al., 2022). Here, as Steller (Steller, 1751) remarked, they used fingerless, bristle-covered forelimbs for support and to shear “*algae and seagrasses from the rocks*”, which they masticated “*not with teeth, which they lack altogether, but with*” large, ridged keratinized pads located on the upper palate and lower mandible. Although they became visibly thin during winter when “*their spinous processes can be seen*”, Steller (Steller, 1751) noted that “*(t)hese animals are very voracious, and eat incessantly*” such that their stupendous stomach (“*6 feet* [1.8 m] *long, 5 feet* [1.5 m] *wide*”) and enormous intestines—which measured a remarkable 5,958 inches (∼151.5 m) from esophagus to anus, equivalent to “*20½ times as long as the whole animal*”—are constantly “*stuffed with food and seaweed*”. The proportionally larger gut (Domning, 2022) is consistent with Steller’s sea cow’s higher energetic requirements relative to extant manatees, which, owing to their low metabolic intensities become cold stressed and die if chronically exposed to water temperatures below 15°C (O’Shea et al., 1985). The inferred reduction in insulative blubber thickness of *H. gigas* during the winter months would likely have compounded the rate of heat loss to sub-zero degree Celsius air and water, though may have been compensated for by arteriovenous anastomoses that regulated blood flow to the skin, and by countercurrent rete supplying the flippers and tail flukes, the latter of which are well developed in manatees and presumably other sirenians (Marshall et al., 2022; Rommel and Caplan, 2003). These structures conserve thermal energy by promoting profound cooling at the appendages and periphery (McCabe et al., 1978), and presumably underlie the low thermal dependence of Steller’s sea cow Hb relative to those of extant sea cows.

Reductions in blood–O_2_ affinity accompanying the *H. gigas* β/δ82Lys→Asn substitution is expected to have further augmented tissue O_2_ delivery without tangible effects on lung O_2_ uptake, thereby helping to fuel thermogenesis and maintain a stable core temperature. In the absence of UCP1-dependent nonshivering thermogenesis (Gaudry et al., 2017), the latter was presumably supplemented by a substantive heat increment arising from fermentation and other post-prandial processes (Marshall et al., 2022). Although the attendant increase in the rate of O_2_ consumption would have mandated a reduction in breath-hold endurance—likely reflecting the relatively short submergence times (4 to 5 minutes) observed by Steller (Steller, 1751)—our results suggest that this may have been partially counteracted by an elevated blood–O_2_ carrying capacity that was potentially coupled to a greater lung volume (Domning, 2022). Underwater foraging times were presumably further defended by key components of the dive reflex, namely bradycardia and peripheral vasoconstriction. Indeed, Steller inadvertently was the first to (indirectly) describe this phenomenon as he observed his crew hunting the animals with spears and knives, “*the blood from the wounded back spurted up like a fountain. As long as he kept his head under water the blood did not flow out, but as soon as he raised his head to breathe the blood leaped forth anew*”.

A final compelling aspect of Steller’s sea cow evolution was their immense size—up to 11,000 kg in mass and 10 m in length—relative to extant sirenians (Domning, 1976). While Steller does not provide measurements of “*their tender little offspring*”, Gerhard Friedrich Müller, who edited Steller’s manuscript prior to publication, noted calves “*weighed 1200 pounds* [544 kg] *and upwards*” (Mueller, 1761). This value is ∼10 to 50 times the mass of new born manatees and dugongs (∼10-50 kg) (Odell, 2009) and is suggestive of rapid prenatal growth during the ∼1 year gestational period indicated by Steller (Steller, 1751). While placental morphology and relative blood flow are important factors affecting pre-natal growth rates, the efficiency of maternal/pre-natal gas exchange is also influenced by differences in blood O_2_-affinity between the two circulations (Carter, 2015). During the early stages of mammalian development, O_2_ diffusion is optimized via the expression of embryonic Hb isoforms with high O_2_-affinity (Weber et al., 1987). Briefly, the α and β gene families of mammals possess multiple paralogs, with the 5’-3’ linkage order and their distance from the respective upstream locus control regions dictating the expression pattern of each locus throughout development (Peterson and Stamatoyannopoulos, 1993). Thus, at two weeks post-conception, developing human embryos begin expressing genes at the 5’ end of the α (HBZ) and β (HBE) clusters, which are translated into ζ- and ε-globin chains, respectively, to form Hb Gower I (Fantoni et al., 1981). At week four, expression of the downstream HBA and HBG loci add α- and γ-chains to the erythrocytes of the developing circulatory system to generate additional Hb isoforms including HbF (α_2_γ_2_) (Hecht et al., 1966). Notably, this pattern of gene expression switching during development results in the temporal production of Hb isoforms with successively lower O_2_ affinities (i.e., each Hb isoform has a lower O_2_ affinity than the protein it replaced), which facilitates O_2_ transfer from maternal to embryonic and fetal blood (Carter, 2015).

In all mammalian lineages examined to date, with the exception of bovid artiodactyls (e.g., goats, sheep, and cows) and simian primates, the expression of the above Hb isoforms is thought to be limited to the embryonic stage of development (Carter, 2015); as such, most mammals express the same Hb isoform (HbA) during both the fetal and post-natal stages of development. However, observations suggest that sea cows and proboscideans (elephants) may also express distinct fetal isoHbs. For example, blood from a 5-month old elephant fetus was shown to contain two distinct Hb components, although only a single (adult) Hb component is present in 12-month old fetal and adult blood (Riegel et al., 1967). Likewise, the blood of manatee calves contains a second isoHb (comprising ∼5% of total Hb; Farmer et al., 1979), which moreover appears to exhibit O_2_ binding properties distinct from that of maternal blood (White et al., 1976). It is thus conceivable that the second Hb component in newborn manatee blood (and potentially other sirenians) arises from the delayed expression of HBG (which expresses the γ-chain of HbF). Given that the timing of globin gene expression is determined by its distance from the locus control region (Peterson and Stamatoyannopoulos, 1993), the possible attenuation of HBG expression in sirenians HBG relative to elephants is supported by synteny comparisons of the β-globin gene cluster, as the HBG locus of sirenians is further downstream than the same locus is in the elephant cluster (see Fig. 1B of ref 4). If expression of the sirenian HBG locus is developmentally delayed to form a discrete isoHb in fetal blood, it would be expected to display P_50_ and cooperativity (n_50_) values that fall between Gower I and HbA, and a response to pH that is similar to the latter.

To test this hypothesis and better understand the maternal/pre-natal gas exchange strategy of sirenians, we thus expressed recombinant Hbs corresponding to Steller’s sea cow Gower I and HbF and dugong HbF (whose γ-chain differs from *H. gigas* γ at four positions (Signore et al., 2019); Fig. S6), and measured their O_2_ binding properties and response to allosteric effectors. As expected, the P_50_ of *H. gigas* Gower I in the combined presence of Cl^-^ and DPG is markedly less than adult Steller’s sea cow Hb (3.1 vs. 15.2 mm Hg, respectively) (Fig. 4A). Similarly, the Hb– O_2_ affinity of Steller’s sea cow and dugong HbF (P_50_ of 0.56 and 1.2 mm Hg, respectively) are substantially higher than that of their respective adult counterparts and, unexpectedly, also higher than that of Steller’s sea cow Gower I (Fig. 4A). In line with the embryonic expressed Hb isoforms of other mammals (Brittain, 2002; Weber et al., 1987), the Bohr and cooperativity coefficients of the sirenian Gower I and HbF proteins were also substantially lower than that of the post-natal (HbA) isoform (Fig. 4A, Table S2). Accordingly, their functional properties are consistent with the embryonic (but not fetal) Hbs of other mammalian species. Although it remains possible that expression of these isoforms lingers into late fetal development, the upstream HBG transcriptional control motif (‘CACCC’) crucial for suppression of human HBB gene expression during the fetal stage (Perez-Stable and Costantini, 1990) is mutated in both dugongs and Steller’s sea cows (but not manatees or elephants; Fig. S7). Consequently, the primary (if not sole) Hb isoform expressed within the both the fetal and post-natal circulation of dugongids is almost certainly HbA. Intriguingly, however, Steller’s sea cow HbF exhibits a distinctly higher O_2_ affinity but lower cooperativity than dugong HbF, traits that are likely attributed to two exceedingly rare γ-chain amino acid replacements positioned within the interior of the protein (γ38Thr→Ile and γ101Glu→Asp; Figs. 4B-E and S6). Briefly, the central cavity γ101Glu→Asp replacement alters the highly conserved sliding interface between the α_1_γ_2_ dimer subunits by forming a hydrogen bond with γ104Arg (Fig. 4B) and has been shown to increase the intrinsic affinity of human Hb Potomac (β101Glu→Asp) (Charache et al., 1978; Shih et al., 1985). Residue γ38 is also potentially functionally relevant as it is located along the α_2_γ_1_ sliding interface and is in contact the distal heme (Fig. 4D) (Ropero et al., 2006). Of note, however, the mRNA capping site of *H. gigas* HbF exhibits an A→G transversion mutation (Fig. S7) that has been shown to lower transcript levels of human HBB by twofold (Meyers et al., 1986). It thus remains unknown if *H. gigas* HbF exhibited a similar downregulation and hence to what degree these γ-chain replacements may have altered O_2_ transfer to the Steller’s sea cow embryo through the amniotic fluid prior to placental development.

**Figure 4.**
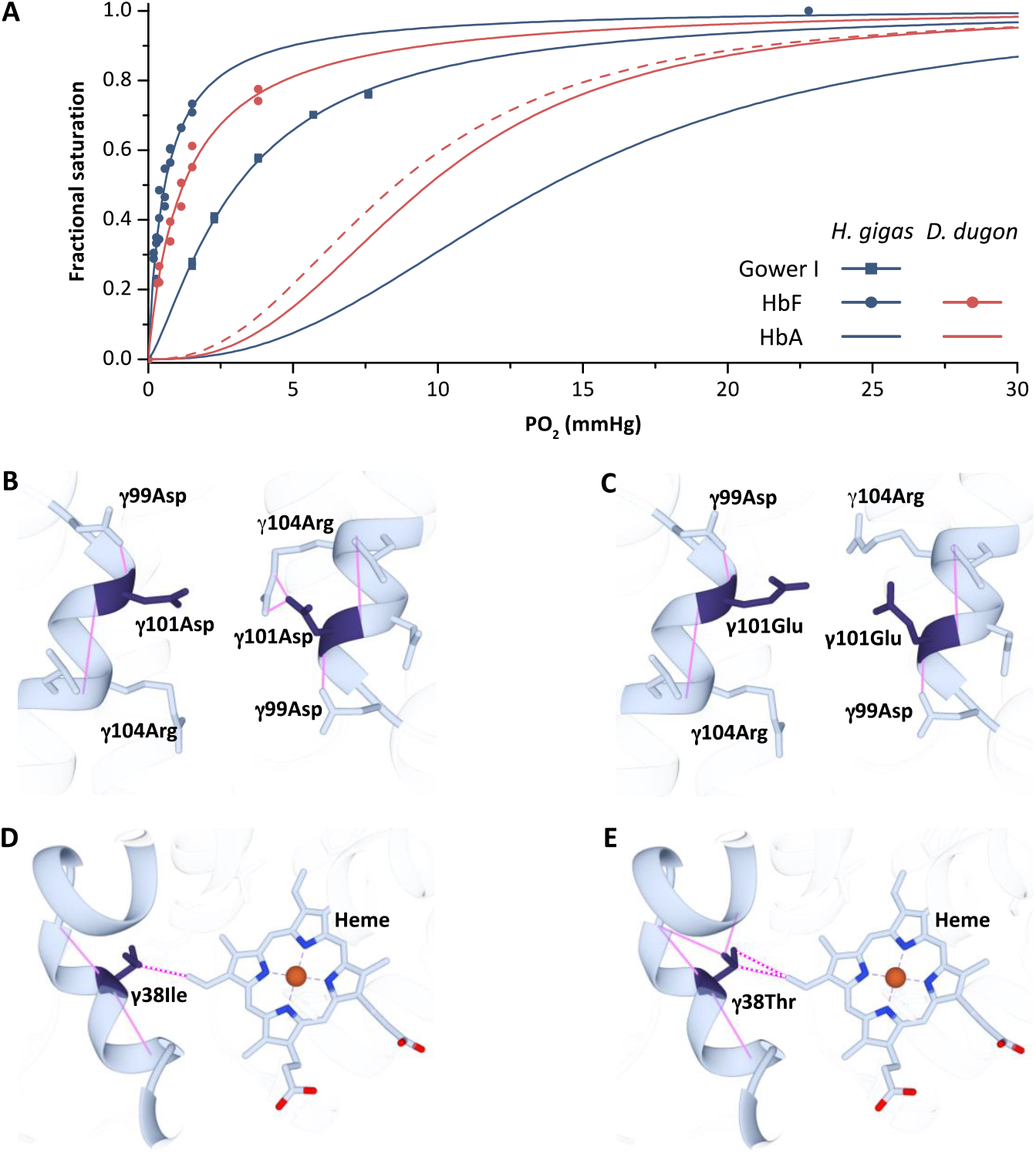
Oxygen equilibrium curves of prenatal and adult sirenian hemoglobins. A) Oxygen equilibrium curves for Steller’s sea cow (*Hydrodamalis gigas*; blue) Hb Gower 1 (ζ_2_ε_2_), HbF (α_2_γ_2_), and HbA (α_2_β/δ_2_) and dugong (*Dugong dugon*; red) HbF and HbA in the presence of allosteric cofactors 2,3-diphosphoglycerate (2-fold molar excess) and KCl (100 mM) at pH 7.1 (prenatal Hbs) or 7.2 (HbA). The dashed red line is for dugong HbA in the absence of DPG that illustrates relative differences in O_2_ affinity between the maternal (solid line) and fetal (dashed line) circulations. Homology models of Steller’s sea cow and dugong HbF denote structural alterations arising from the *H. gigas* specific γ101 (B vs. C, respectively) and γ38 (D vs. E, respectively) replacements. Solid pink lines denote predicted hydrogen bonds while the dashed pink lines represent predicted Van der Waals interactions.

Based on the above considerations, the apparently unique DPG insensitive phenotype of *H. gigas* HbA is particularly noteworthy owing to its potential impact on maternal/fetal O_2_ exchange. Presumably to assist in this process, fetal blood cells expressing HbA contain only trace amounts of DPG hence conferring fetal blood with an elevated O_2_ affinity relative to that of the maternal circulation (compare the dashed vs. solid red lines in Fig. 4A as an example) in the vast majority of mammalian species (Bunn, 1980; Carter, 2015). By contrast, and owing to the inability of Steller’s sea cows Hb to respond to DPG in either the fetal or adult circulations, this species would represent a rare example (feloids and eastern moles are others) in which fetal and maternal blood have the same (albeit relatively low) O_2_ affinity. However, placental O_2_ delivery in these species will be defended by the well-known double Bohr effect, whereby CO_2_ transport from the fetal to maternal circulation increases blood O_2_ affinity in the former while lowering it in the latter (Carter, 2015). More importantly, the evolved reduction in blood O_2_ affinity of Steller’s sea cows would have stipulated that equilibrium between umbilical and uterine blood was reached at a higher PO_2_, a condition that is expected to substantially improve O_2_ delivery to the fetal circulation (Carter, 2015). The lower fetal blood O_2_ affinity (relative to manatees and dugongs) and concomitant higher fetal blood-to-tissue PO_2_ gradients are further expected to have augmented O_2_ delivery to the developing tissues of this species. As such, these attributes, together with increases in Hb solubility/reduced susceptibility to oxidative damage arising from β/δ82Lys→Asn that conceivably also elevated the O_2_ carrying capacity of fetal blood, may have been important contributors to the enhanced fetal growth rate of these immense sirenians. The resulting increase in thermal inertia and relatively low surface-area-to-volume ratio following birth, together with an adaptively reduced Hb thermal sensitivity and thick ‘bark-like’ skin arising from inactivation of lipoxygenase genes (Le Duc et al., 2022), were presumably central components of Steller’s sea cows successful exploitation of the harsh sub-Arctic marine environments of the North Pacific.

## Materials and Methods

### Sequence Collection and Analyses

The pre-natal (HBZ-T1, HBE, and HBG) and adult-expressed Hb genes (HBA and HBB/HBD) of the Florida manatee, dugong, and Steller’s sea cow, and the most recent common ancestor shared by Steller’s sea cow and the dugong (‘ancestral dugongid’) have previously been determined (Signore et al., 2019). As the *H. gigas* β/δ82Lys→Asn exchange is not known to occur in any living species, we mined recently deposited genomes for 13 additional Steller’s sea cows (PRJNA484555, PRJEB43951) to test for the prevalence of this replacement in the population. Briefly, we first searched SRA files of each specimen using the megablast function against a previously determined *H. gigas HBB/HBD* gene sequence (GenBank accession #: MK562081). All hits were then downloaded, trimmed of adapters and low-quality regions using BBDuk (Joint Genome Institute), and assembled to *H. gigas HBB/HBD* using Geneious Prime 2019 software (Biomatters Ltd, Auckland, New Zealand). Assemblies generated using genome reads that were not pre-treated with uracil-DNA glycosylase and endonuclease VIII to reduce C→T and G→A damage artifacts (Le Duc et al., 2022) were examined to ensure these deamination artifacts did not affect the consensus sequences.

### Construction of Recombinant Hb Expression Vectors

Coding sequences for Steller’s sea cow Gower I (ζ_2_ε_2_), dugong and H. gigas HbF (α_2_γ_2_), and the above four HbA (α_2_β/δ_2_) proteins were optimized for expression in *E. coli* and synthesized *in vitro* by GenScript (Piscataway, NJ). The resulting gene cassettes were digested with restriction enzymes and tandemly ligated into a custom Hb expression vector (Natarajan et al., 2011) using a New England BioLabs Quick Ligation Kit as recommended by the manufacturer. Chemically competent JM109 (DE3) *E. coli* (Promega) were prepared using a Z-Competent *E. coli* Transformation Kit and Buffer Set (Zymo Research). We also prepared a *H. gigas* β/δ82Asn→Lys Hb mutant via site-directed mutagenesis on the Steller’s sea cow Hb expression vector by whole plasmid amplification using mutagenic primers and Phusion High-Fidelity DNA Polymerase (New England BioLabs), phosphorylation with T4 Polynucleotide Kinase (New England BioLabs), and circularization with an NEB Quick Ligation Kit (New England BioLabs). All site-directed mutagenesis steps were performed using the manufacture’s recommended protocol.

Hb expression vectors were co-transformed into JM109 (DE3) chemically competent *E. coli* alongside a plasmid expressing methionine aminopeptidase (Natarajan et al., 2011), plated on LB agar containing ampicillin (100 μg/ml) and kanamycin (50 μg/ml), and incubated for 16 hours at 37**°**C. A single colony from each transformation was cultured in 50 ml of 2xYT broth for 16 hours at 37**°**C while shaking at 200 rpm. Post incubation, 5 ml of the culture was pelleted by centrifugation and plasmid DNA was isolated using a GeneJET Plasmid Miniprep Kit (Thermo Scientific). The plasmid sequence was verified using BigDye 3.1 sequencing chemistry and an ABI3130 Genetic Analyzer. The remainder of the culture was supplemented with glycerol to a final concentration of 10%, divided into 25 ml aliquots and stored at -80**°**C until needed for expression.

### Expression and Purification of Recombinant Hb

25 ml of starter culture (above) was added to 1250 ml of TB media containing ampicillin (100 μg/ul) and kanamycin (50 μg/ul) and distributed evenly amongst five 1 L Erlenmeyer flasks. Cultures were grown at 37**°**C while shaking at 200 rpm until the absorbance at 600 nm reached 0.6-0.8. Hb expression was induced by supplementing the media with 0.2 mM isopropyl β-D-1-thiogalactopyranoside, 50 μg/ml of hemin and 20 g/L of glucose and the culture was incubated at 28**°**C for 16 hours while shaking at 200 rpm. Once expression had completed, dissolved O_2_ was removed by adding sodium dithionite (1 mg/ml) to the culture, which was promptly saturated with CO for 15 minutes. Bacterial cells were then pelleted by centrifugation and Hb purified by ion exchange chromatography according to Natarajan et al. (Natarajan et al., 2011)

It should be noted that the β82Asn residue of human Hb Providence is relatively uncommon in that it slowly undergoes post-translational deamidation *in vivo* to form aspartic acid, with the latter residue (β82Asp) comprising ∼67-75% in mature mixed blood (Bardakjian et al., 1985; Perutz et al., 1980). While it is unknown to what degree *H. gigas* β/δ82Asn was catalyzed into Asp in nature, O_2_ binding data (see below) of this species was collected from freshly purified recombinant samples for which only one peak—presumably β/δ82Asn—was resolved during chromatography (data not shown). Additionally, this reaction is dependent on the local protein environment (Robinson, 2002), specifically two nearby residues β143His and β83Gly (Perutz et al., 1980). Importantly, the latter residue was replaced by β/δ83Ser on the Steller’s sea cow branch (Figs. 1B and 3B), which is expected to slow (but not stop) the rate of deamidation (Robinson, 2002). Regardless, since the two Hb Providence isoforms have similar O_2_ affinities and functional properties (Bardakjian et al., 1985; Bonaventura et al., 1976; Charache et al., 1977) it is unlikely that presence of β/δ82Asp in Steller’s sea cow blood would meaningfully alter the results and interpretations presented herein.

### Functional Analyses of Hbs

O_2_-equillibrium curves for HbA containing solutions (0.25–1.0 mM heme in 0.1 M HEPES/0.0005 M EDTA buffers) were measured at 25 and 37**°**C using the thin film technique (Weber, 1992), while curves for the three pre-natal Hb isoforms (0.25 mM heme in 0.1 M HEPES/0.0005 M EDTA) were measured at 37**°**C using a multi-cuvette tonometer cell described by Lilly et al. (Lilly et al., 2013). Hb solutions varied in their pH (range: 6.8 to 7.9), chloride concentration (0 or 0.1 M KCl), and organic phosphate concentration (0 or 2-fold molar excess of DPG relative to tetrameric Hb concentrations) in order to test the influence of these cofactors on Hb function. Each Hb solution was sequentially equilibrated with gas mixtures of three to five different oxygen tensions (PO_2_) that result in Hb–O_2_ saturations between 30 to 70%. Hill plots (log[fractional saturation/[1-fractional saturation]] vs logPO_2_) constructed from these measurements were used to determine the PO_2_ (P_50_) and the cooperativity coefficient (n_50_) at half saturation, from the χ-intercept and slope of these plots, respectively. By this method, the *r*^2^ determination coefficients for the fitted curves exceed 0.995 and the standard errors (SEM) are less than 3% of the P_50_ and n_50_ values (Weber et al., 2014). A linear regression was fit to plots of log*P*_50_ vs. pH, and the resulting equation was used to estimate *P*_50_, Cl^-^ effect, and DPG effect values (± SE of the regression estimate) at pH 7.20 for HbA samples, and pH 7.10 for Gower I and HbF samples (to account for the lower pH of pre-natal blood). The slope of these plots (ΔlogP_50_/ΔpH) represented the Bohr effect. P_50_ values at 25 and 37**°**C were used to assess the thermal sensitivity of the Hbs by calculating the apparent enthalpy of oxygenation using the van’t Hoff isochore:

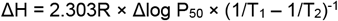

where R is the universal gas constant and T_1_ and T_2_ are the absolute temperatures (**°**K) at which the P_50_ values were measured. All ΔH values were corrected for the heat of O_2_ solubilization (12.55 kJ mol^-1^ O_2_).

### Solubility assay

Ammonium sulfate was added to Hb solutions (0.074±0.004 mM Hb_4_) to generate final concentrations that ranged from 0 to 3.5 M. These solutions were incubated for 60 minutes at 37°C and the remaining soluble Hb was measured via Drabkin’s reagent, according to the manufacturer’s instructions (Sigma-Aldrich).

### Homology modelling

To assess the structural effect of the *H. gigas* specific β/δ replacements on the DPG binding site, homology models of ancestral dugongid and Steller’s sea cow Hb were constructed using the SWISS-MODEL server (62) using the three-dimensional human deoxy structure with DPG bound (PDB: 1B86) as a template (30). The sequence conservation of amino acid residues implicated in DPG binding were calculated by the ConSurf Server (63) from a subsample of 51 mammalian beta-type hemoglobin chains downloaded from GenBank (Table S3). Homology models were visualized with UCSF Chimera (64). To assess the structural differences between *H. gigas* and *D. dugon* HbF (α_2_γ_2_), homology models of these proteins were created as above, but with human deoxy HbF used as template (PDB: 4MQJ).

## Acknowledgments

This manuscript is dedicated to the memory of our dear friend Joseph (Joe) Bonaventura for his pioneering work on the human Hb Providence Asn/Asp proteins. We thank Mike Gaudry, Diana Hanna, and Elin Ellebæk Petersen for technical assistance, Chandrasekhar Natarajan for providing us with a hemoglobin expression plasmid, and Jay Storz for constructive feedback on an earlier version of this manuscript. Authorization to use paintings by Carl Buell was kindly provided by John Gatesy. This study was supported by NSERC (Canada) Discovery and Accelerator Supplement Grants (RGPIN/238838-2011, RGPIN/412336-2011, and RGPIN/06562-2016 to K.L.C; RGPIN/261924-2013 and RGPIN/446005-2013 to C.J.B.), an NSERC Postgraduate Scholarship (A.V.S.), the Faculty of Science and Technology, Aarhus University (R.E.W.), and the Independent Research Fund Denmark (A.F.; Danmarks Frie Forskningsråd DFF-4181-00094).

