## Supplemental Information for "Evolution of an extreme hemoglobin phenotype contributed to the sub-Arctic specialization of extinct Steller’s sea cows"

**This PDF file includes:**

Figures S1 to S7
Tables S1 to S3

**HBA**

**Exon 1**

```
               1        10        20        30
T. manatus     VLSDEDKTNVKTFWGKIGTHTGEYGGEALER
Anc. dugongid  ...A............L.A..A.........
D. dugon       ...A............L.A..A...S.....
H. gigas       ...A............L.A..A.........
```

**Exon 2**

```
                       40        50        60        70        80        90
T. manatus     MFLSFPTTKTYFPHFDLSHGSGQIKAHGKKVADALTRAVGHLEDLPGTLSELSDLHAHRLRVDPVNFK
Anc. dugongid  ................M....D....................D.....................I...
D. dugon       ..NA..A.........M....D..........E.........D.......D.............I...
H. gigas       ................MK.D.D....................D...............K.....I...
```

**Exon 3**

```
               100       110       120       130       140
T. manatus     LLSHCLLVTLSSHLREDFTPSVHASLDKFLSSVSTVLTSKYR
Anc. dugongid  ..............P.....P..........N..........
D. dugon       ...........N..PD....P..........N..........
H. gigas       ...........G..P.....P..........N..........
```

**HBB/HBD**

**Exon 1**

```
               1        10        20        30
T. manatus     VHLTPEEKALVIGLWAKVNVKEYGGEALGR
Anc. dugongid  ....AD.....T..................
D. dugon       ....AD.T...T..................
H. gigas       ....AD.....T...S..............
```

**Exon 2**

```
                       40        50        60        70        80        90       100
T. manatus     LLVVYPWTQRFFEHFGDLSSASAIMNNPKVKAHGEKVFTSFGDGLKHLEDLKGAFAELSELHCDKLHVDPENFR
Anc. dugongid  .......................V.H...........LA.........D.........................
D. dugon       .......................V.H...........LA.........D..........A...E......Q..K
H. gigas       .......................V.H.S..QT.....LA.........DN.NS.....................
```

**Exon 3**

```
                   110       120       130       140
T. manatus     LLGNVLVCVLARHFGKEFSPEAQAAYQKVVAGVANALAHKYH
Anc. dugongid  .............L......Q.....................
D. dugon       ....M.....S..L......Q.....E...............
H. gigas       .............L......Q.....................
```

**Fig. S1.** Amino acid sequences of sirenian *HBA* and *HBB/HBD* genes and the reconstructed sequences of the last common ancestor (‘Anc. dugongid’) shared by the dugong (*Dugong dugon*) and Steller’s sea cow (*Hydrodamalis gigas*). Dots represent sequence identity with the Florida manatee (*Trichechus manatus latirostris*).

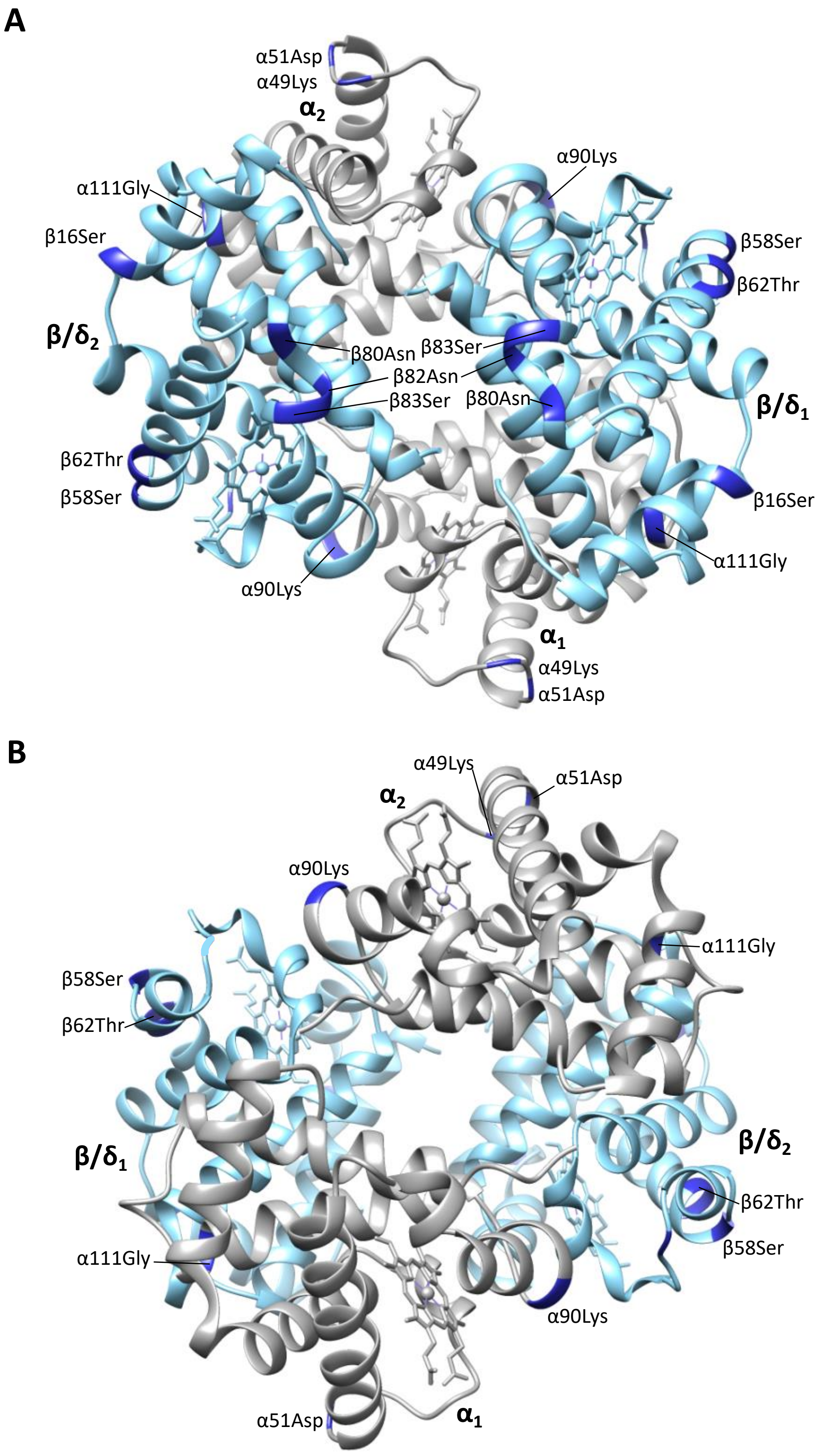


**Fig. S2.** Homology model of Steller's sea cow (*Hydrodamalis gigas*) adult-expressed ($\alpha_2\beta/\delta_2$) hemoglobin with A) the β/δ-subunits (blue) in the foreground and B) the α-subunits (grey) in the foreground. Dark blue colored residues represent the 11 *H. gigas* specific substitutions.

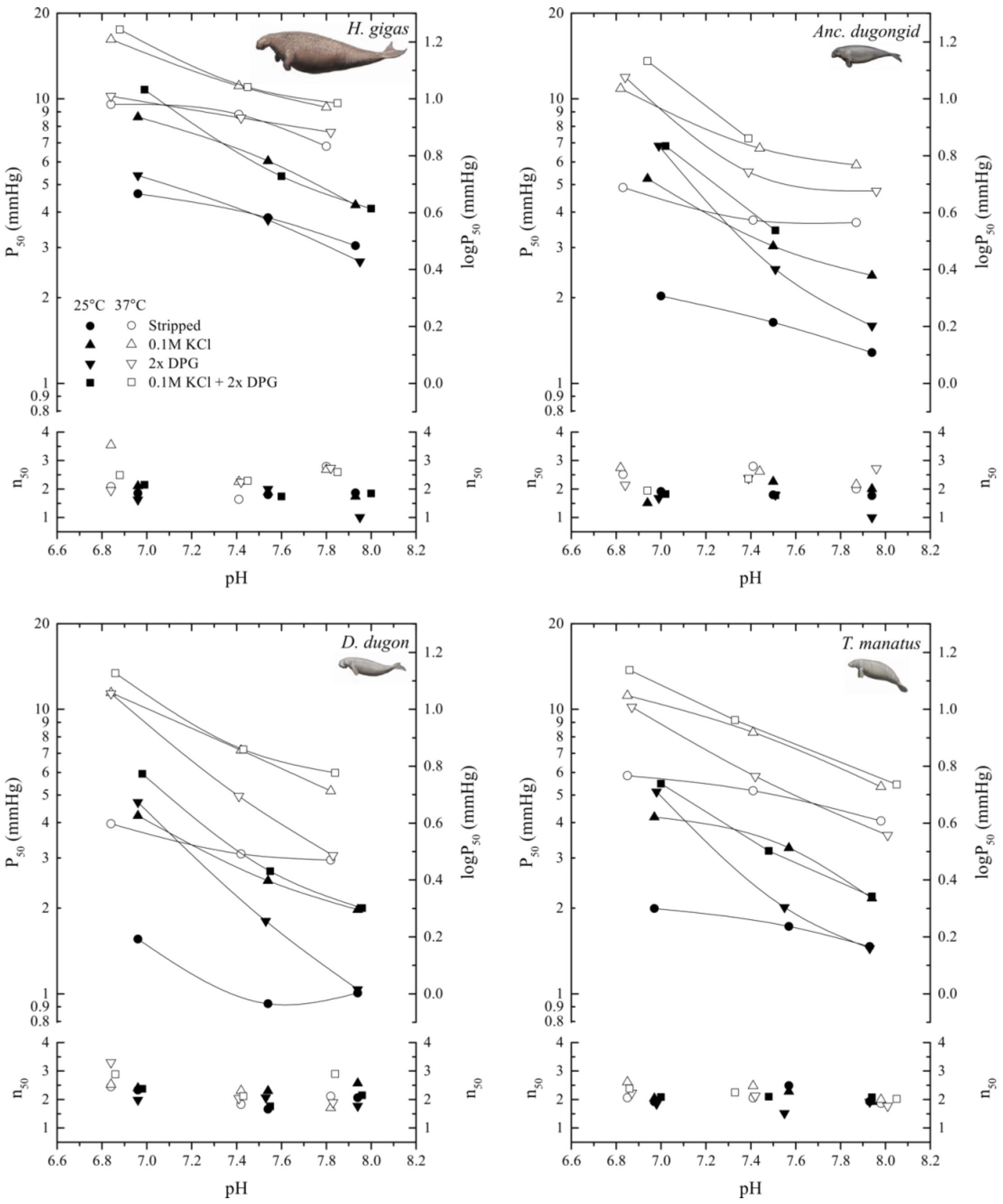


**Fig. S3.** The pH dependence of oxygen tensions and the cooperativity coefficients at half $O_2$ saturation ($P_{50}$ and $n_{50}$, respectively) for hemoglobins of the Florida manatee (*Trichechus manatus latirostris*), dugong (*Dugong dugon*), Steller's sea cow (*Hydrodamalis gigas*), and the last common dugongid ancestor ('Anc. dugongid') in stripped Hb (circles), and in the presence of 0.1 M KCl (triangles), of a 2-fold molar excess of 2,3-diphosphoglycerate (DPG; inverted triangles), and of both KCl and DPG (squares), at 25°C (solid symbols) and 37°C (open symbols).

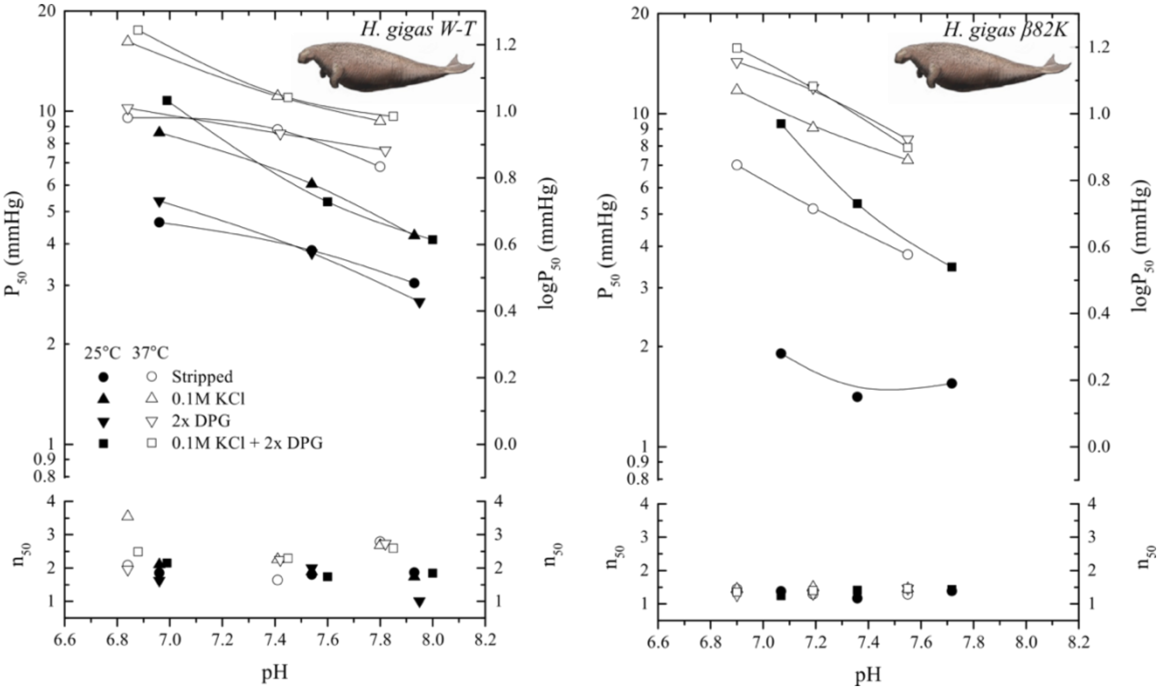


**Fig. S4.** The pH dependence of oxygen tensions and the cooperativity coefficients at half $O_2$ saturation ($P_{50}$ and $n_{50}$, respectively) of wild-type Steller's sea cow hemoglobin (*H.gigas W-T*) and a mutated Steller's sea cow β/δ82Lys variant (*H. gigas β82K*) in the absence and presence of allosteric effectors (0.1 M KCl and/or 2-fold molar excess of 2,3-diphosphoglycerate (DPG)) at 25°C (solid symbols) and 37°C (open symbols).

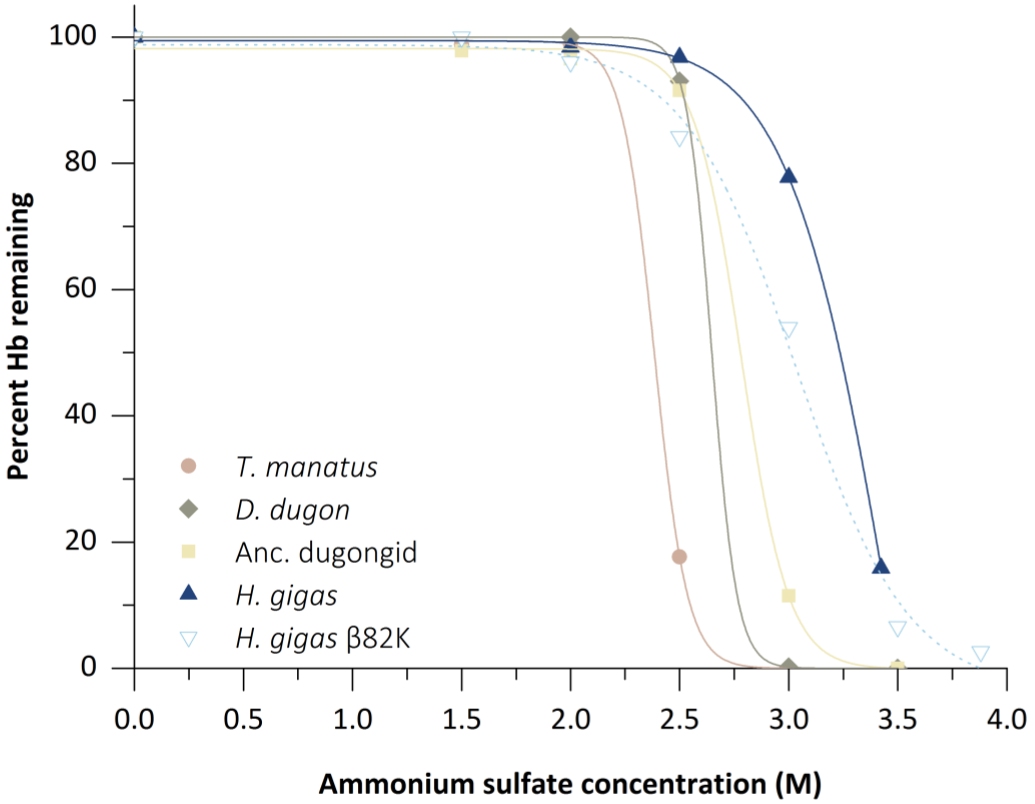


**Fig. S5.** Solubility assay of five sirenian hemoglobins (Hb) illustrating the percentage of Hb protein remaining in solution after precipitation by the addition of ammonium sulfate.

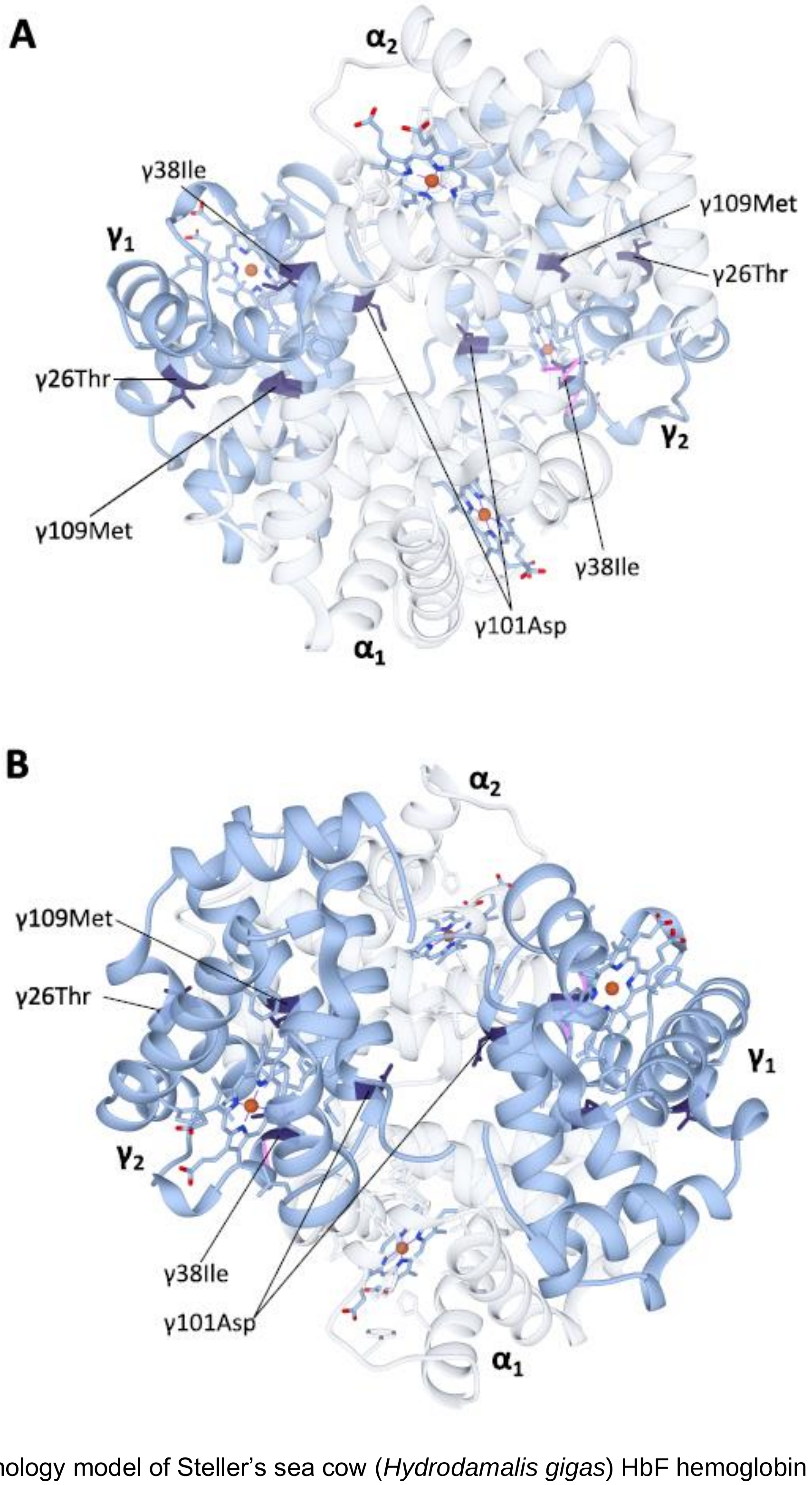


**Fig. S6.** Homology model of Steller's sea cow (*Hydrodamalis gigas*) HbF hemoglobin ($\alpha_2\gamma_2$) with A) the $\alpha$-subunits in the foreground and B) the $\gamma$-subunits in the foreground. The $\alpha$-subunits are identical to those in the *H. gigas* adult-expressed hemoglobin (see Fig. S4 for details) and have been made transparent to improve clarity of the $\gamma$-subunit substitutions. Dark blue colored residues denote the four *H. gigas* specific $\gamma$-chain substitutions ($\gamma$26Lys→Thr, $\gamma$38Thr→Ile, $\gamma$101Glu→Asp, and $\gamma$109Val→Met) identified by Signore et al. (2019).

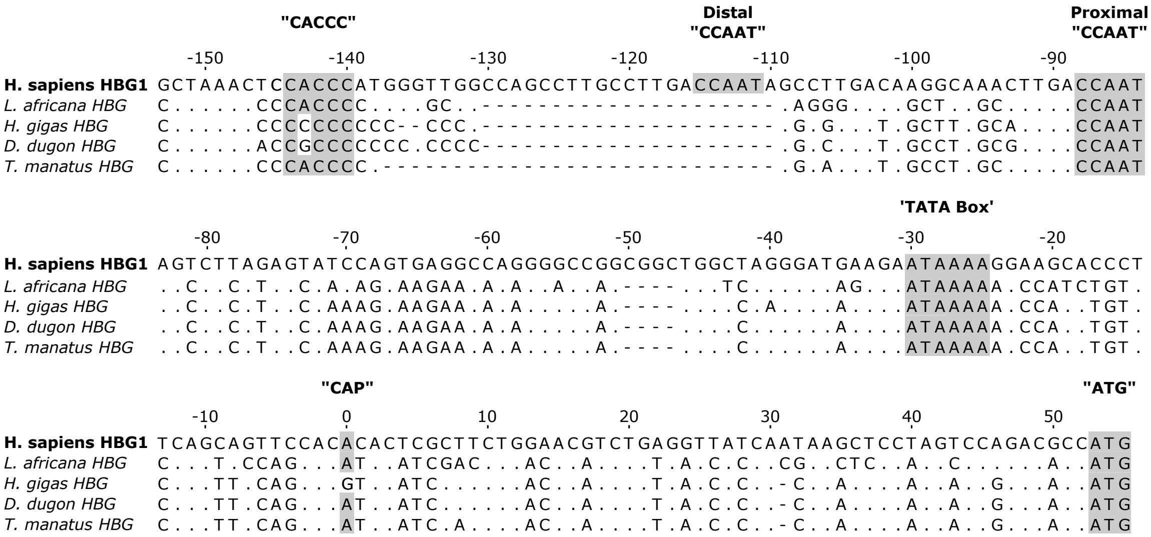


**Fig. S7.** DNA alignment of the 5' untranslated region and transcriptional promoter regions for representative paenungulate and human HBG genes (hyrax HBG is pseudogenized (Signore et al. 2019) and hence not included here). Regulatory elements and the "ATG" start codons are highlighted in grey. Dots represent sequence identity with *Homo sapiens*, while hyphens represent alignment gaps. Note that the "CACCC" promoter element required for fetal suppression of HBB is mutated in both Steller's sea cows (*Hydrodamalis gigas*) and dugongs (*Dugong dugon*). Conversely, this box element is intact in both the African elephant (*Loxodonta africana*) and Florida manatee (*Trichechus manatus latirostris*) HbF promoter region, consistent with a fetal expression pattern in these species. Note also the A→G mutation at the putative mRNA "CAP" site of *H. gigas* HBG that has been shown to downregulate the level of human HBB transcription by approximately two-fold (Meyers et al., 1986).

**Table S1.** Intrinsic oxygen affinities ($P_{50}$, mmHg) for the adult-expressed hemoglobins ($\alpha_2\beta/\delta_2$) of the Florida manatee (*Trichechus manatus latirostris*), dugong (*Dugong dugon*), Steller's sea cow (*Hydrodamalis gigas*), and the ancestral dugongid ('Anc. dugongid'), and their sensitivity to allosteric effectors at 25 and 37°C in 0.1 M HEPES buffer. All values are corrected to pH 7.2.

| Species | Temp | [a]$P_{50}$ | [a]$\log P_{50}$ | [b]$Cl^-$ effect | [c]DPG effect | [d]Bohr effect ($\Delta \log P_{50}/\Delta pH$) | | | |
|---|---|---|---|---|---|---|---|---|---|
| | | | | | | Stripped | +KCl | +DPG | KCl + DPG |
| *T. manatus* | 37°C | 5.36 | 0.73 | 0.23 | 0.14 | -0.14 | -0.28 | -0.39 | -0.34 |
| | 25°C | 1.88 | 0.27 | 0.30 | 0.29 | -0.14 | -0.28 | -0.59 | -0.42 |
| *Anc. dugongid* | 37°C | 4.30 | 0.63 | 0.29 | 0.27 | -0.12 | -0.26 | -0.36 | -0.60 |
| | 25°C | 1.86 | 0.27 | 0.35 | 0.40 | -0.21 | -0.34 | -0.67 | -0.61 |
| *D. dugon* | 37°C | 3.51 | 0.55 | 0.39 | 0.30 | -0.13 | -0.35 | -0.58 | -0.36 |
| | 25°C | 1.29 | 0.11 | 0.42 | 0.39 | -0.21 | -0.34 | -0.68 | -0.49 |
| *H. gigas* | 37°C | 8.80 | 0.94 | 0.17 | 0.02 | -0.15 | -0.25 | -0.13 | -0.27 |
| | 25°C | 4.25 | 0.63 | 0.24 | 0.03 | -0.19 | -0.32 | -0.30 | -0.42 |
| *H. gigas* β82k | 37°C | 5.39 | 0.71 | 0.25 | 0.27 | -0.41 | -0.32 | -0.36 | -0.46 |
| | 25°C | 1.72 | 0.23 | – | – | -0.14 | – | – | -0.65 |

[a]$PO_2$ at half-saturation (mm Hg)

[b]$\Delta \log P_{50}^{(KCl - stripped)}$

[c]$\Delta \log P_{50}^{(DPG - stripped)}$

[d]pH range 6.9 to 7.8

**Table S2.** Intrinsic oxygen affinities ($P_{50}$, mmHg) for the prenatally-expressed hemoglobins Gower I ($\zeta_2\varepsilon_2$) and HbF ($\alpha_2\gamma_2$) of Steller's sea cow (*Hydrodamalis gigas*) and HbF ($\alpha_2\gamma_2$) of the dugong (*Dugong dugon*), and their sensitivity to allosteric effectors at 37°C in 0.1 M HEPES buffer. All values are corrected to pH 7.1 to account for the lower pH of prenatal blood.

| Hb | [a]$P_{50}$ | [a]$\log P_{50}$ | [b]$Cl^-$ effect | [c]DPG effect | [d]Bohr effect ($\Delta \log P_{50}/\Delta pH$) | | | |
|---|---|---|---|---|---|---|---|---|
| | | | | | Stripped | +KCl | +DPG | KCl + DPG |
| *H. gigas* Gower I | 1.90 | 0.28 | 0.07 | 0.23 | 0.18 | -0.05 | -0.21 | 0.20 |
| *H. gigas* HbF | 0.53 | -0.27 | -0.15 | 0.13 | 0.07 | -0.50 | 0.58 | -0.09 |
| *D. dugon* HbF | 0.57 | -0.24 | 0.40 | 0.20 | -0.10 | 0.68 | 0.56 | 0.16 |

[a]$PO_2$ at half-saturation (mm Hg)

[b]$\Delta \log P_{50}^{(KCl - stripped)}$

[c]$\Delta \log P_{50}^{(DPG - stripped)}$

[d]pH range 6.75 to 7.15

**Table S3.** Accession numbers for beta-globin amino acid sequences used for ConSurf analyses.

| Organism | Common name | Accession # |
|---|---|---|
| *Alces alces alces* | European elk | P02073 |
| *Ammotragus lervia* | Aoudad | ABC86528 |
| *Ateles geoffroyi* | Black-handed spider monkey | P68232 |
| *Balaenoptera acutorostrata* | Minke whale | P18984 |
| *Bison bonasus* | European bison | P09422 |
| *Bos taurus* | Cattle | P02070 |
| *Bradypus tridactylus* | Pale-throated sloth | AAZ22685 |
| *Bubalus bubalis* | Water buffalo | P67820 |
| *Camelus bactrianus* | Bactrian camel | P68230 |
| *Canis lupus familiaris* | Domestic Dog | P60524 |
| *Capra hircus* | Goat | P02077 |
| *Cavia porcellus* | Domestic guinea pig | P02095 |
| *Ceratotherium simum* | White rhinoceros | P02066 |
| *Dasypus novemcinctus* | Nine-banded armadillo | AAZ22684 |
| *Didelphis virginiana* | North American opossum | P02109 |
| *Dugong dugon* | Dugong | QBK14998 |
| *Echinops telfairi* | Small Madagascar hedgehog | AAB21591 |
| *Elephas maximus* | Asiatic elephant | ACV41403 |
| *Equus caballus* | Horse | P02062 |
| *Erinaceus europaeus* | Western European hedgehog | P02059 |
| *Eulemur fulvus fulvus* | Common brown lemur | P02053 |
| *Felis catus* | Domestic cat | P07412 |
| *Hippopotamus amphibius* | Hippopotamus | P19016 |
| *Homo sapiens* | Human | P68871 |
| *Lama glama* | Llama | P68226 |
| *Loxodonta africana* | African savanna elephant | ACV41399 |
| *Macaca mulatta* | Rhesus monkey | P02026 |
| *Macropus eugenii* | Tammar wallaby | Q6H1U7 |
| *Macropus giganteus* | Eastern gray kangaroo | P02106 |
| *Mammuthus primigenius* | Woolly mammoth | ACV41408 |
| *Myotis velifer* | Mouse-eared bat | P11758 |
| *Odocoileus virginianus virginianus* | Virginia white-tailed deer | P02074 |
| *Ornithorhynchus anatinus* | Platypus | P02111 |
| *Oryctolagus cuniculus* | Rabbit | P02057 |
| *Ovis aries musimon* | European mouflon | P02076 |
| *Procavia capensis* | Cape rock hyrax | Q45XI6 |
| *Rangifer tarandus* | Reindeer | P21380 |
| *Rattus norvegicus* | Norway rat | P02091 |
| *Rousettus aegyptiacus* | Egyptian rousette | P02058 |
| *Suncus murinus* | House shrew | P02060 |
| *Sus scrofa* | Pig | P02067 |
| *Tachyglossus aculeatus aculeatus* | Short-beaked echidna | P02110 |
| *Talpa europaea* | European mole | P02061 |
| *Tapirus terrestris* | Brazilian tapir | P02064 |
| *Tarsius bancanus* | Horsfield's tarsier | P02051 |
| *Tragelaphus strepsiceros* | Greater kudu | P04245 |
| *Trichechus manatus* | West Indian manatee | Q45XI8 |
| *Tupaia glis* | Common tree shrew | P02052 |
| *Tursiops truncatus* | Bottlenose dolphin | P18990 |
| *Ursus maritimus* | Polar bear | P68011 |
| *Varecia variegata* | Ruffed lemur | P21667 |
